# Input dose differentiation by NF-κB

**DOI:** 10.1101/752394

**Authors:** Minjun Son, Andrew Wang, Hsiung-Lin Tu, Marie O Metzig, Parthiv Patel, Kabir Husain, Jing Lin, Arvind Murugan, Alexander Hoffmann, Savaş Tay

## Abstract

Cells receive a wide range of dynamic signaling inputs during immune regulation, but how gene regulatory networks measure and interpret such dynamic inputs is not understood. Here, we used microfluidic live-cell analysis and mathematical modeling to study how NF-κB pathway in single-cells responds to time-varying immune inputs such as increasing, decreasing or fluctuating cytokine signals. Surprisingly, we found that NF-κB acts as a differentiator, responding strictly to the absolute difference in cytokine concentration, and not to the concentration itself. Our analyses revealed that negative feedbacks by the regulatory proteins A20 and IκBα enable dose differentiation by providing short-term memory of prior cytokine level and continuously resetting kinase cycling and receptor levels. Investigation of NF-κB target gene expression showed that cells create unique transcriptional responses under different dynamic cytokine profiles. Our results demonstrate how cells use simple network motifs and transcription factor dynamics to efficiently extract information from complex signaling environments.

## Introduction

Cells receive temporally varying signals like cytokines and pathogenic molecules that encode information on the identity, amplitude and timing of extracellular challenges during the immune response. This information is transmitted into the nuclear localization dynamics of key transcription factors like NF-κB, which is then decoded by gene regulatory machinery to create specific gene expression responses^1–4^. Among those signaling molecules are the central inflammatory cytokines TNF and IL-1β, which mediate local activation of neighboring cells and coordinate population and system-level responses. In response to infection in humans and mice, TNF and IL-1β levels in tissue and plasma show a rapid increase and can be sustained at a high level for hours up to many days^5–10^. On the other hand, the cytokine levels in the tissue microenvironment can be highly dynamic^11–13^. Individual macrophages and T-cells secrete cytokines in complex temporal profiles such as pulses and oscillations in an input specific manner^14–16^. These and other observations led to the hypothesis that cytokine dynamics and the subsequent pro-inflammatory responses are connected by a “temporal code” that transmits information from cell surface receptors to gene expresion^17^.

NF-κB is a key mediator of cytokine signaling and is central to a wide range of physiological scenarios in immunity and disease, such as infection, autoimmunity and cancer. Nuclear translocation of NF-κB family transcription factors like p65 leads to the activation of nearly one thousand inflammatory and immune response genes^18^. Both population averaged and single-cell resolved measurements show that NF-κB activation discriminates between different doses of signaling molecules like TNF and LPS^4,19–21^. However, it is not known whether cells perceive and respond to the absolute concentration of the cytokine signals independent of past cytokine exposure, as it would be for a memory-less system, or to the relative changes in cytokine levels where cells have memory of the previous cytokine levels. Additionally, it is not known how gene regulatory networks like NF-κB interpret increasing or decreasing concentrations of extracellular signaling molecules, i.e., whether they respond to the absolute cytokine level or to the rate of change.

To address this important question, we investigated how single mammalian cells respond to dynamically changing cytokine concentrations. Using high-throughput microfluidic live cell analysis, single cell tracking, and mathematical modeling, we investigated whether the NF-κB system exploits the temporal characteristics of signals (i.e. the input dynamics) during inflammatory signaling (Fig. 1C). Surprisingly, we find that single cell NF-κB dynamics precisely measures the rate of change in TNF and IL-1β concentration. This intriguing behavior is analogous to a differentiation operation seen in many engineering systems, but has not previously been described in biological systems. Our theoretical and experimental analyses revealed that the multipoint negative feedbacks by A20 and IκBα proteins enable this dose differentiation behavior in individual cells, and that NF-κB target genes effectively distinguish different dynamic signaling patterns suggesting a functional role for dose differentiation in NF-κB function.

**Figure 1.**
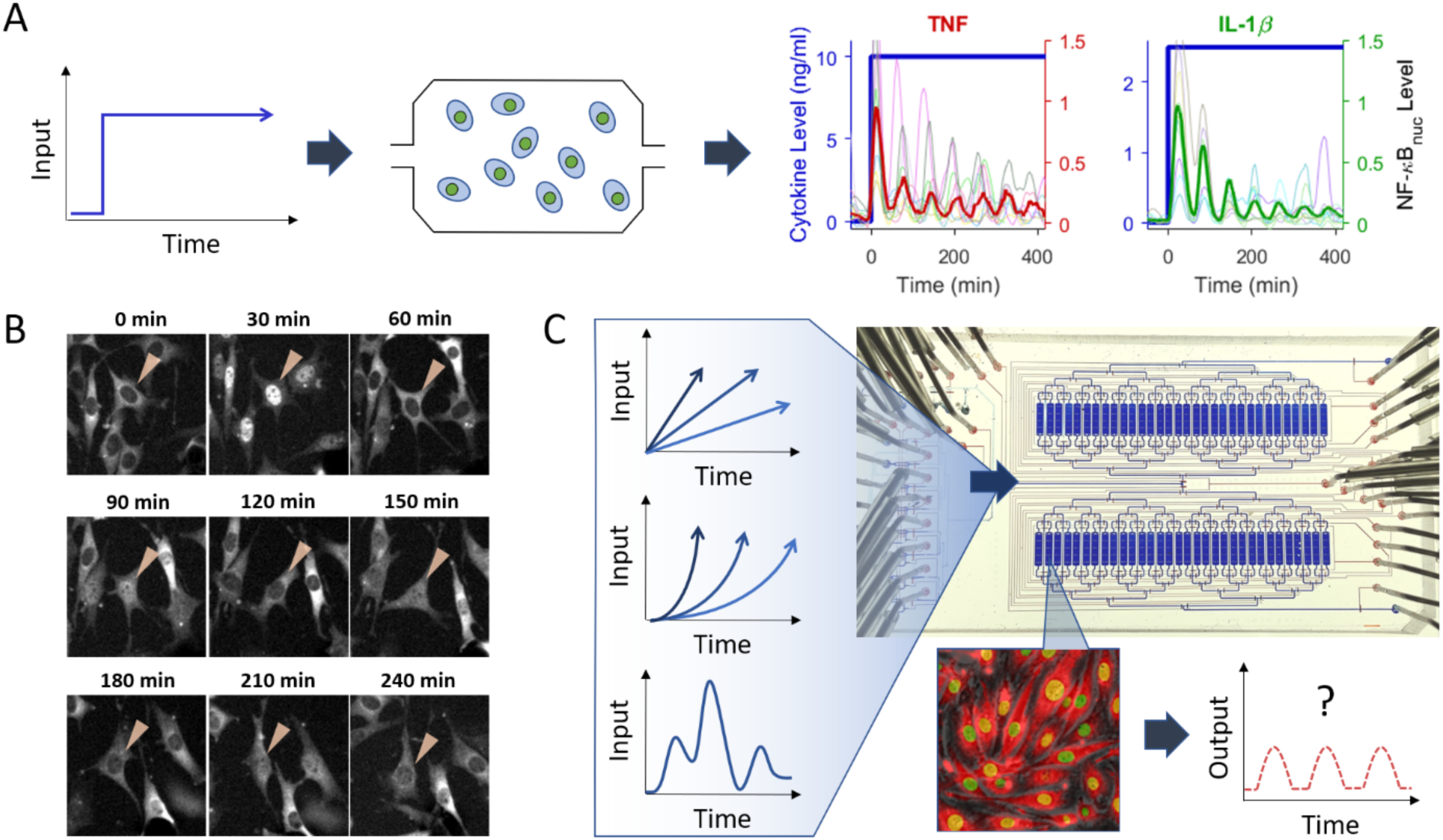
Single live-cell analysis enables the study of NF-κB response to dynamical signaling inputs. (A, B) Cells were exposed to instantly increased high dose of TNF (red) or IL-1β (green) in a microfluidic device. The thick red or green line indicates the median of single cell nuclear NF-κB traces, while the thin colored lines represent examples of measured single cell traces (n >200 for each condition). NF-κB nuclear localization traces show an initial strong response followed by sharp decay, as shown in the representative NF-κB fluorescence time-series in (B). (C) To test how NF-κB responds to different input dynamics, cytokine levels (designated as ‘Input’) were increased linearly or exponentially with various rates or doubling times, or were introduced in a randomly fluctuating order. Dynamic inputs were introduced to cells expressing p65-DsRed and H2B-GFP, through a multiplexed microfluidic cell culture chip. Nuclear NF-κB localization was measured by time-lapse microscopy.

## Results

### NF-κB precisely measures the rate of change in TNF and IL-1β concentration

To understand how dynamically varying cytokine concentrations are encoded and decoded by a signaling system, two key questions need to be answered: which attributes of the dynamic signals are measured by the responding pathway and how these attributes regulate the response dynamics. The characteristic response of NF-κB transcription factors to cytokine stimulus is oscillatory. The transcription factor p65 moves into the nucleus and cycles between the cytosol and nucleus upon cellular exposure to cytokine signals^1,4^ (Fig. 1A). These oscillations can be measured by live-cell microscopy using fluorescent reporters^4^. We stimulated 3T3 mouse embryonic fibroblasts (MEFs) expressing the p65-DsRed fusion protein with various dynamic cytokine regiments created by a microfluidic device, and analyzed peak height and area-under-curve (AUC) of each p65 (NF-κB) localization peak, the two measures of total NF-κB nuclear localization closely correlated with downstream gene expression^4,22–24^.

To study which input detection mechanism is employed by NF-κB, we delivered periodically increasing doses of TNF and IL-1β to 3T3 fibroblasts in a microfluidic culture device. Extracellular concentration of both signaling molecules were increased in three different patterns (instant, linear, and exponential) in a precisely controlled manner (Fig. 1C). Individual cells were tracked in real-time, and nuclear p65-DsRed levels were recorded via video microscopy. Computer-aided analysis of the videos allowed extraction of nuclear p65 localization traces for each cell.

When TNF or IL-1β was instantly increased to a high dose (10 or 2.5 ng/ml) and maintained at that level, a strong initial nuclear translocation of NF-κB was observed in all cells within 5 min, reaching its peak at around 25 min (Fig. 1A, Supplementary Movie 1 and 2). However, even though the dose was maintained at the same level, the amplitude of subsequent nuclear NF-κB oscillations was diminished (10 – 30% of the initial peak height). The fact that the NF-κB response exponentially decayed under constant cytokine level indicates that NF-κB system desensitizes its response to the unchanging input.

When TNF or IL-1β was linearly increased to a high dose (1 or 0.5 ng/ml increment every 60 min), we observed strong sustained oscillations (Fig. 2A, also see Supplementary Movie 1 and 2 for cell images). The p65 peaks under both TNF and IL-1β linear ramping increased and were sustained at a high level until ramping was terminated. The total p65 localization, measured by AUC of nuclear p65 during each ramping step, stayed constant under the linearly increasing cytokine dose. After seeing clear differences in oscillation characteristics between instantaneous increase and linear step-wise ramping, we studied the NF-κB response under exponentially increasing concentration steps. The cytokine concentration was doubled every 60 min until reaching the maximum level. During exponential ramping, we observed that p65 peaks increased continuously until the final concentration was reached (Fig. 2B, see Supplementary Movie 1 and 2 for cell images). Remarkably, when plotted against the logarithm of input change, the mean of AUC in each step fitted precisely to a straight line (adjusted R-squared value ∼ 0.963).

**Figure 2.**
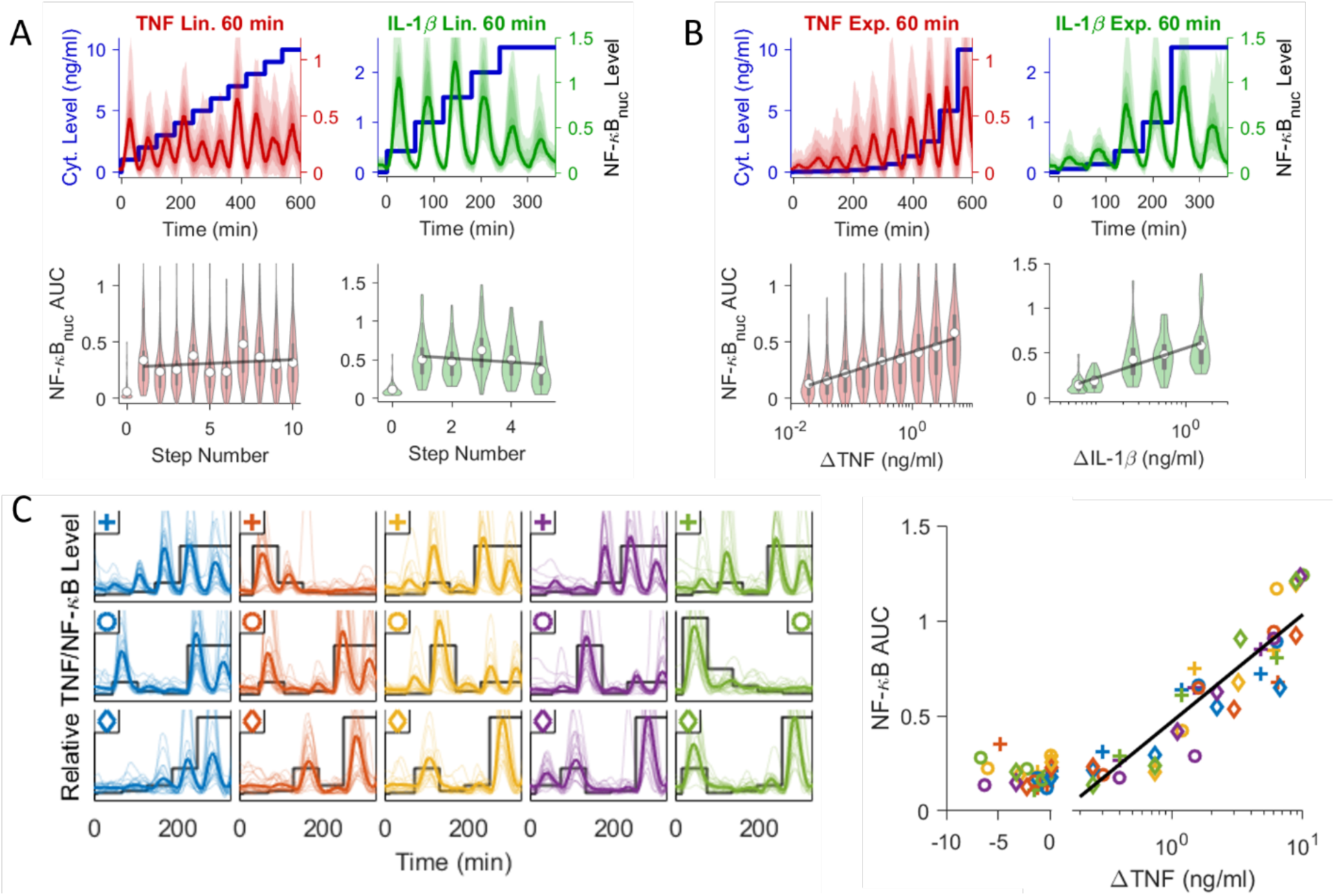
Dose differentiation by the NF-κB pathway. (A, B) TNF and IL-1β concentrations were increased linearly or exponentially and nuclear NF-κB levels were recorded from individual cells. The thick blue line shows the time-dependent cytokine concentration, and thick red and green lines show median nuclear NF-κB level during linear (A) and exponential (B) input ramping. The color gradient indicates the range of single cell response, with darkest being the 40^th^ to 60^th^ percentile and lightest being the 10^th^ or 90^th^ percentile. The bottom violin plots show the distribution of NF-κB area under curve (AUC) at each cytokine dose step, where the white circles represent the mean of single cell AUCs and the best fit based on differentiator behavior is shown by the thick black thick lines. (C) Cells were exposed to variable TNF concentrations in random order. The black line shows TNF input dose ranging from 0 to 10 ng/ml, the thick colored line shows the mean NF-κB trace (n ∼ 200), and the thin lines show 20 random single cell traces for each sample. On the right scatter plot, the AUC after each dose increase/decrease was calculated and plotted in linear scale for negative ΔTNF (TNF concentration change), and log scale for positive ΔTNF. The black line indicates the best fit to the log function.

Our data showed constant NF-κB peak heights and AUCs under linear ramping, and increasing peak heights and AUCs under exponential ramping. This behavior can be best described as a dose “differentiator” (red output line in Fig. 1C), where NF-κB is responding to the log-difference in cytokine dose (and not to its absolute level or the fold-change), similar to an electronic circuit whose output is proportional to the rate of change of the input level.

Having observed NF-κB behaving as a dose differentiator under specific cytokine ramping conditions, we asked how robustly the differentiation behavior holds across variable input dynamics. To investigate how the NF-κB system processes randomly fluctuating cytokine levels, we stimulated cells with different levels of TNF/IL-1β in randomly increasing and decreasing concentrations every 60 min (Fig. 2C for TNF and Supplementary Fig. 3C for IL-1β). The AUC of each time interval for each sample was calculated then plotted against the change of the cytokine level at each step. For all random dynamics we tested, any decrease in cytokine level resulted in a uniformly sharp decrease in nuclear AUC. However, when cytokine levels increased, the AUC maintained a linear relationship to the log of the input change (correlation coefficient ∼ 0.91). Our results thus indicate that the NF-κB system robustly differentiates positive rate of change in the TNF/IL-1β dose regardless of input complexity.

Overall, all our results indicate that the NF-κB is analogous to a rectified differentiator, except that its response is proportional to the logarithm of the input difference. We also find that the differentiator behavior in NF-κB holds robustly over a wide range of ramping or exponential growth rates in physiological timescales (0.75 – 10 ng/ml/h or 6 – 120 min doubling time), further supporting the generality of differentiator behavior under various input dynamics (Supplementary Fig. 1, 2, and 3). Thus, the following simple expression best describes the relationship between the input stimulus and nuclear NF-κB localization amplitude or AUC in single cells, for positive changes in cytokine concentrations (Δ*C*_*cytokine*_ > 0):

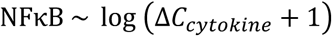

### NF-κB dose differentiation is unique to cytokine signals

To investigate if the differentiator behavior is applicable to the response to the dynamics of other types of ligands, we tested NF-κB response to the dynamics of Pam2CSK4 (Pam), a chemically synthetic diacylated lipopeptide, which mimics bacterial lipoproteins and acts as a TLR2 agonist (Fig. 3)^25^. NF-κB response decreases exponentially when Pam was increased instantaneously, just like it did for TNF and IL-1β. However, when Pam was increased exponentially, the median NF-κB response exhibited sudden increase around 1 ng/ml at early time point, then showed gradual decrease afterwards (Fig. 3A), which was not observed for the exponential TNF and IL-1β increase.

**Figure 3.**
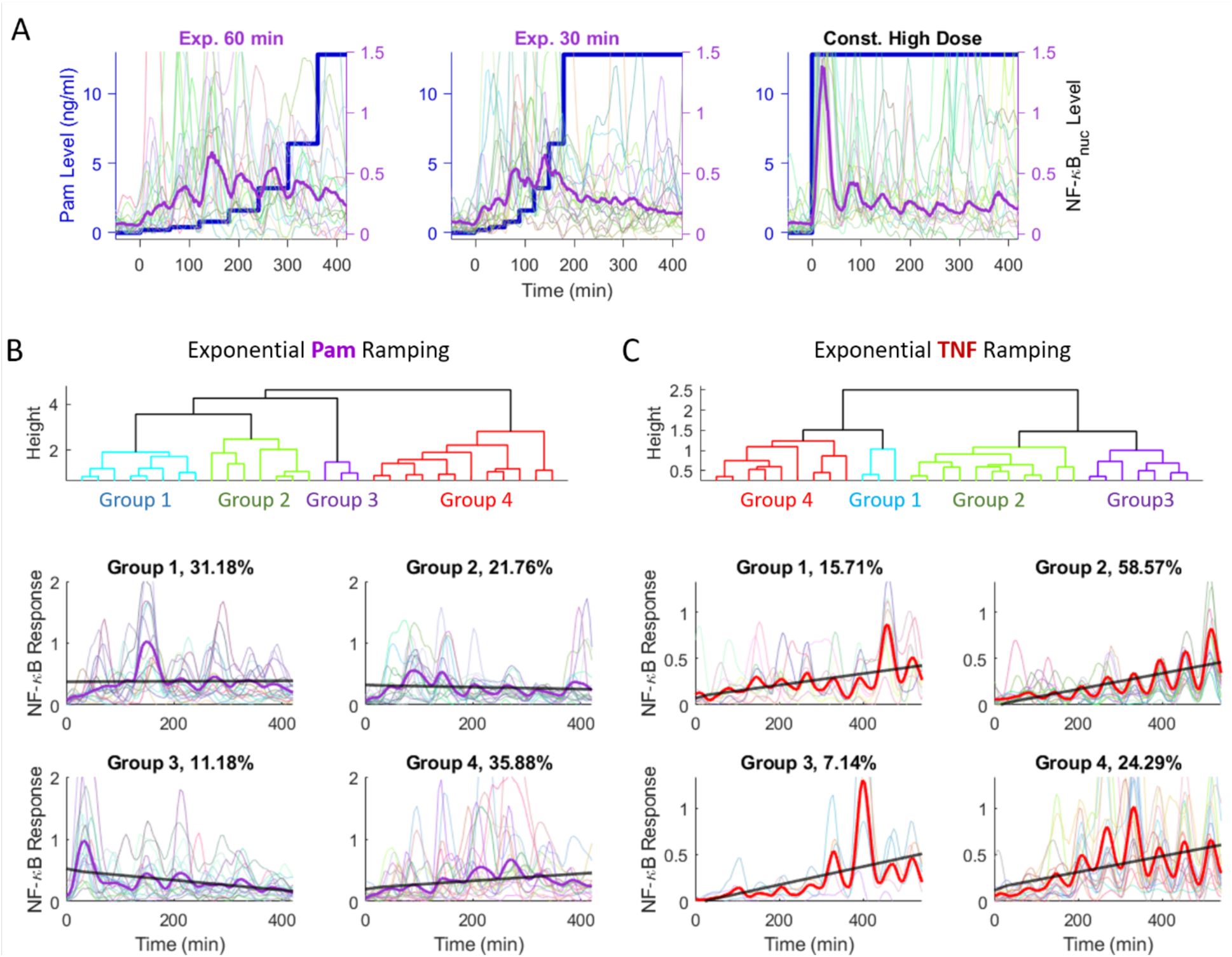
Ramping with Pam3CSK4 shows the input differentiation in the NF-κB system is specific to certain ligand types. (A) For exponential ramping, Pam was doubled every 30 or 60 min from 0.2 to 12.8 ng/ml, while cells were directly exposed to 12.8 ng/ml in the instant increase experiment. Here, the blue lines show the Pam dose, while purple lines indicate the mean trace of nuclear NF-κB. The mean trace appears to resemble TNF/IL-1β ramping, seemingly following the dose increase up to certain level (∼ 1 ng/ml), while cells show fast adaptation (desensitization) to the instant increase. (B, C). However, single cell analysis highlights fundamental differences between the Pam and TNF ramping. The exponential ramping data for both ligands were clustered into four groups using Matlab’s ‘linkage’ function (‘ward’ method and ‘correlation’ metric). The dendrogram in the top shows the clustering from the Pam (B) and TNF (C) exponential ramping data. The bottom subplots show the single cell and mean traces for each cluster, where the numbers in the title indicate the percent of cells belonging to each group. The thick gray line in each group shows the best fit linear line for AUC in each ligand step, similar to the gray lines in Fig. 2A and B.

To examine if this different response to exponential ramping persists at the single cell level, we grouped each trace into four different clusters using ‘linkage’ method through Matlab (Fig. 3B and C). The AUC at each ramping step was calculated for the mean trace and was fitted to linear line (gray thick line in Fig. 3B and C), in order to evaluate the differentiator behavior of each cluster. For Pam stimulation, majority of cells exhibited one or two sharp peaks during the entire ramping period (Fig. 3B). In fact, few cells exhibited robust NF-κB oscillation, and the slope of the fitted line fluctuated from minus to positive value (0.0021, -0.0095, -0.0481, and 0.0349 for Group 1 to 4 respectively). This result indicates NF-κB system does not effectively differentiate the dose change of Pam. On the other hand, the AUC at each TNF ramping step exhibited gradual increase even at the single cell level (Fig. 3C). Although the variability existed due to stochastic activation, most cells exhibited continuous oscillation and generally increasing peak heights. The slope of the fitted line was all positive (0.0373, 0.0515, 0.0585, and 0.0512 for Group 1 to 4 respectively), and all slopes were higher than the highest slope in the Pam clusters. Overall, our single cell analysis indicates that the NF-κB system selectively processes the input dynamics depending on the types of input molecules.

### Dose differentiation depends on IKK dynamics and negative feedback acting upstream of IKK

To understand the molecular mechanism behind dose differentiation, we studied specific properties of the NF-κB regulatory network through mathematical modeling. When not stimulated, cells harbor the majority of their NF-κB in the cytosol, sequestered by IκB proteins (Fig. 4A). Upon activation by TNF or IL-1β ligands, the membrane receptors trigger the ubiquitination and phosphorylation of neutral IκB kinase (IKK_n_), which transforms the kinase into an active conformation (IKK_a_) and cause the degradation of IκB protein. The degradation releases NF-κB for nuclear transport and triggers the transcription of immune response genes including its own negative regulators, IκB and A20. The newly expressed IκB sequesters nuclear NF-κB and exports the complex out to the cytosol. Coincident with NF-κB translocation, IKK_a_ undergoes conformational changes and phosphorylation at different sites that disrupt its catalytic activity, leaving it in a refractory state (IKK_i_)^22,26,27^. Dephosphorylation and reformation of the IKK subunits is necessary in order to adopt the poised state (IKK_n_) again. Theoretical studies demonstrated that the cycling rates of IKK conformational change can affect the peak height and dynamics of NF-κB activity significantly^28^. If IKK_a_ remains at a high level after NF-κB and IκB complex has been exported, it triggers degradation of IκB, initiating another cycle of NF-κB translocation. Another important negative feedback component regulated by nuclear NF-κB is A20, a ubiquitin editing enzyme that inhibits the activation of IKK and facilitates the inactivation of IKK_a_^29–31^. Both *in vitro* and *in vivo* studies showed that defects in A20 cause prolonged nuclear residence of NF-κB and increased immune activity^32,33^.

**Figure 4.**
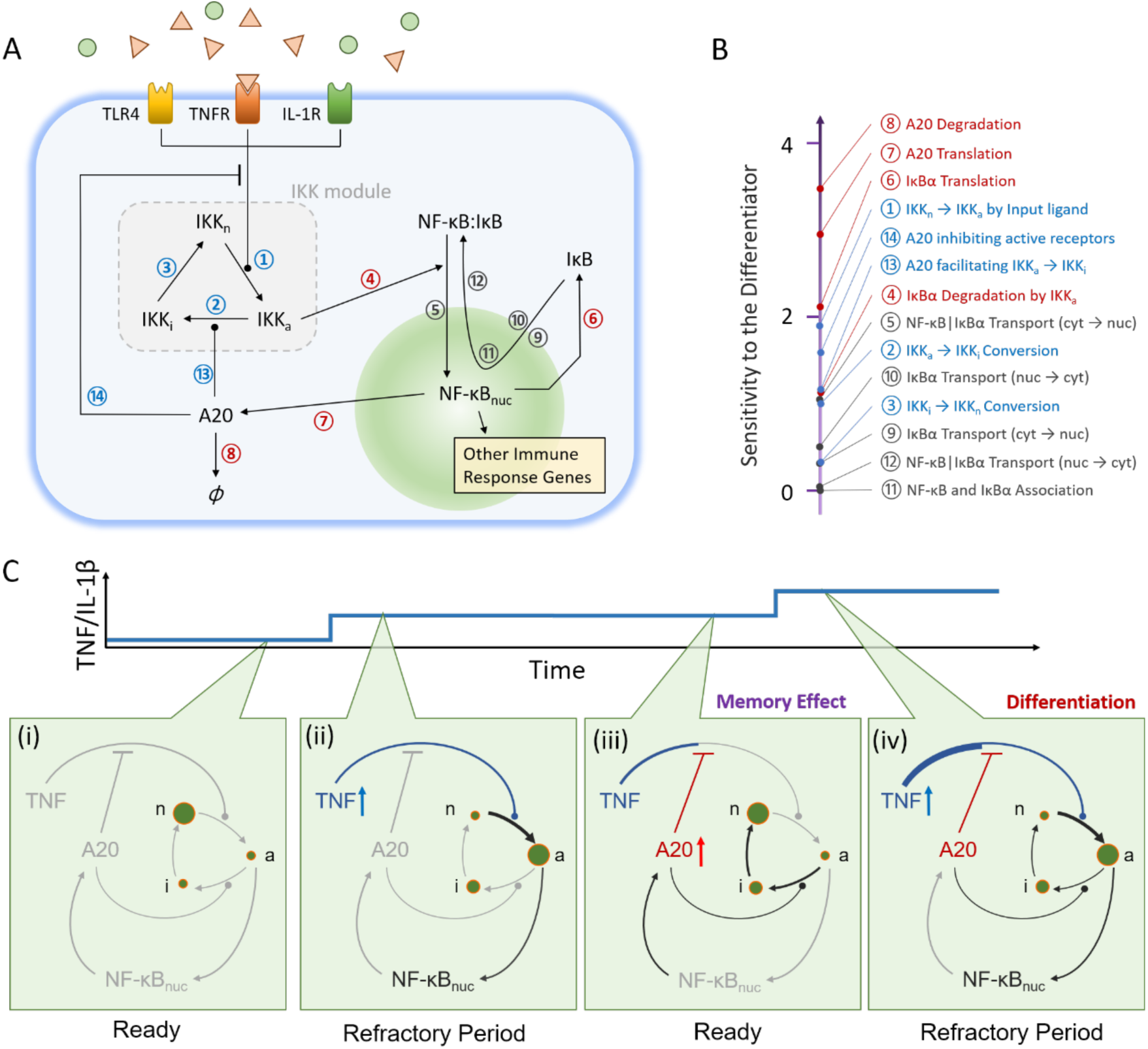
Computational analysis shows that A20 facilitates dose differentiation by providing memory to NF-κB. (A) The NF-κB circuit describes how cytokine input regulates immune response. Parameters governing cycling dynamics of IKK are in blue, parameters associated with production and degradation of negative feedback molecules (IκBα and A20) are in red, and other parameters used in the sensitivity test are in black. (B) For each parameter indicated in diagram (A), the value of the parameter was computationally varied by 10-fold. Then the change in the statistical distance (χ^2^) to the differentiating behavior was calculated (Mathematical Method in Supplementary Information), which was used as sensitivity score. More detailed analysis can be found in Supplementary Fig. 4. (C) Based on the sensitivity test in (B), the schematics of the differentiator mechanism is predicted. (i) Before stimulation with ligand, the system is at the stationary phase with most IKK in the neutral state (IKK_n_). (ii) Introduction of TNF/IL-1β converts neutral IKK into active form (IKK_a_), which triggers the translocation of NF-κB into nucleus. (iii) The translocation initiates the production of A20, which inhibits the receptors activated by the ligand and deactivates the active IKK. Here, the amount or activity of A20 depends on the strength or dose of the previous stimulation; A20 thus serves as “memory” of the previous stimulation by dampening the subsequent response. (iv) However, if the dose of the ligand is increased, more receptors become active overcoming the inhibition by A20.

To computationally identify possible molecular components or NF-κB sub-circuits underlying dose differentiation, we constructed a system of ten differential equations with twenty-five parameters (see Mathematical Method in Supplementary Information for model fitting), based on our previously published models of the NF-κB pathway^4,34^. Approximately half of the parameters could be estimated based on previous experimental work. Then, using the simplex search algorithm^35^, the parameter values for the other variables were estimated by fitting the simulated nuclear NF-κB dynamics under three different input ramping patterns (instant, linear, and exponential increase) to the corresponding experimental data (Fig. 5A). The simulated nuclear NF-κB trajectory closely captured the differentiator trait of our experimental data. Using this parameter set as a basis, we then evaluated the sensitivity of each parameter to the differentiator behavior through a custom developed scoring system (Fig. 4B, Supplementary Fig. 4, Mathematical Method in Supplementary Information).

**Figure 5.**
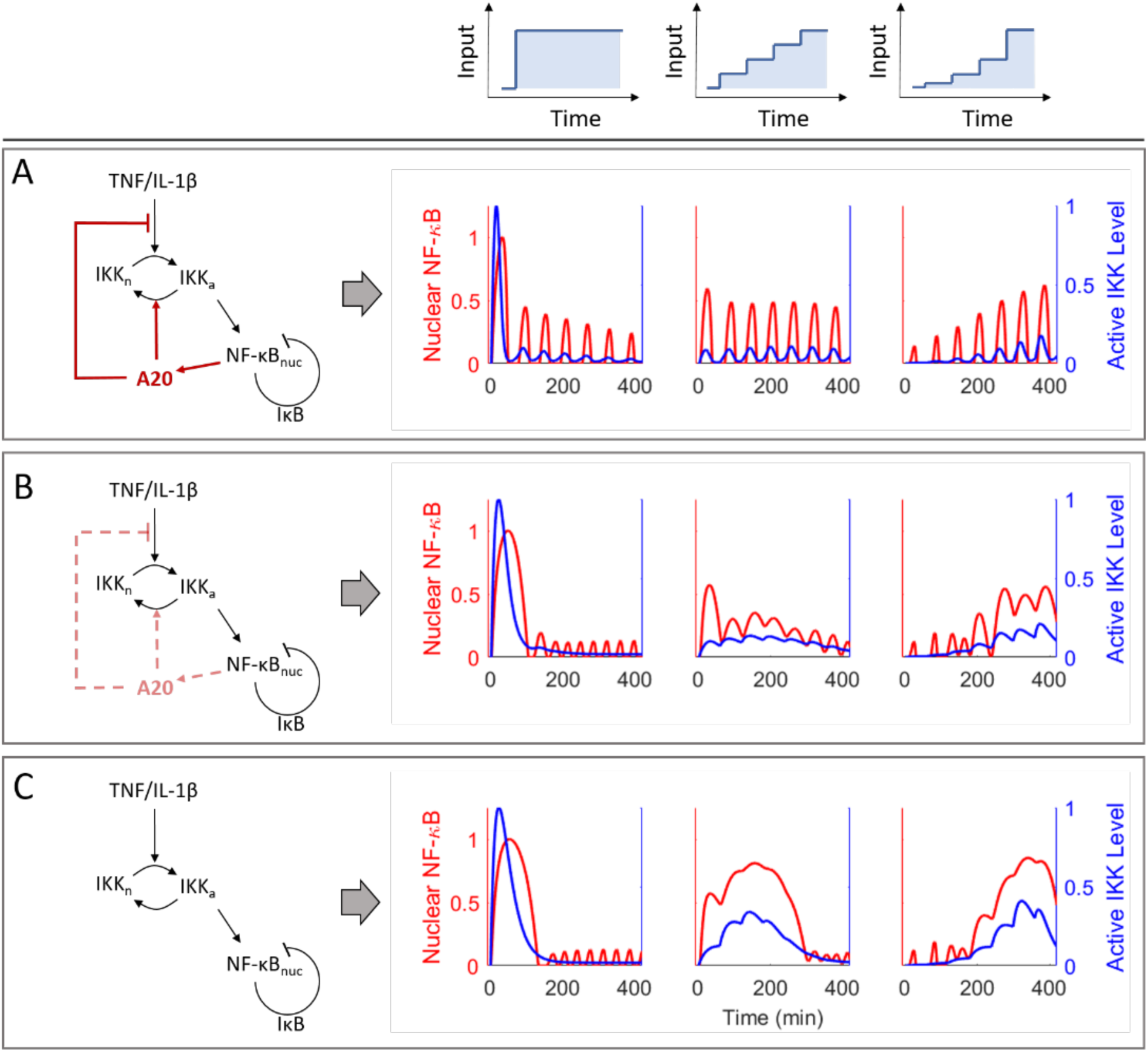
NF-κB network simulation demonstrates the role of A20 in input differentiation. The role of A20 and IKK cycling in the NF-κB differentiator was modeled (A-C). (A) The red and blue curves represent the dynamics of nuclear NF-κB and active IKK levels under three different input dynamics. The red lines in the circuit diagram highlight the A20 negative feedback, which is investigated in this simulation (B, C) The translation rate of A20 is reduced to quarter of the original value (B) or is set to zero (C). The reduced production of A20 undermines the control of upstream signaling through nuclear NF-κB (dotted red line), disrupting the differentiation capability of the system (B). Eliminating negative feedback to upstream components results in greater disruption of the dose differentiation behavior (C).

We found that differentiator behavior was largely unaffected by changes in any of the molecular transport rates (export and import of NF-κB and IκB through nuclear membrane, black parameters in Fig. 4A and B). The rates associated with IKK conformational change (blue parameters) exhibited moderate effects. However, the terms associated with expression of A20 protein showed significant effects when varied (red parameters). The importance of A20 parameters for differentiator behavior could be explained by the dependence of A20 on the strength or dose of the previous TNF stimulation; which would provide memory of the previous TNF stimulation by damping the following NF-κB response. The high sensitivity of the dose differentiation behavior to A20 is intriguing, as this is the only element of the model that is not essential to the canonical NF-κB pathway. To experimentally validate the importance of A20 for differentiator behavior under dynamic input, we deprived cells of A20 via CRISPR/Cas9, which led to a reduction of protein levels by ∼ 90% (Supplementary Fig. 6), and stimulated these cells with exponentially increasing doses of TNF (Fig. 6A). A20 knockout MEFs (A20 KO) elicited sharp increases in both peak and trough heights and caused the response to saturate at a much lower dose of TNF, effectively abolishing the differentiator response. Both our simulation and perturbation experiments highlighted the importance of A20 in the dose differentiator behavior as an emergent property of the NF-κB system.

**Figure 6.**
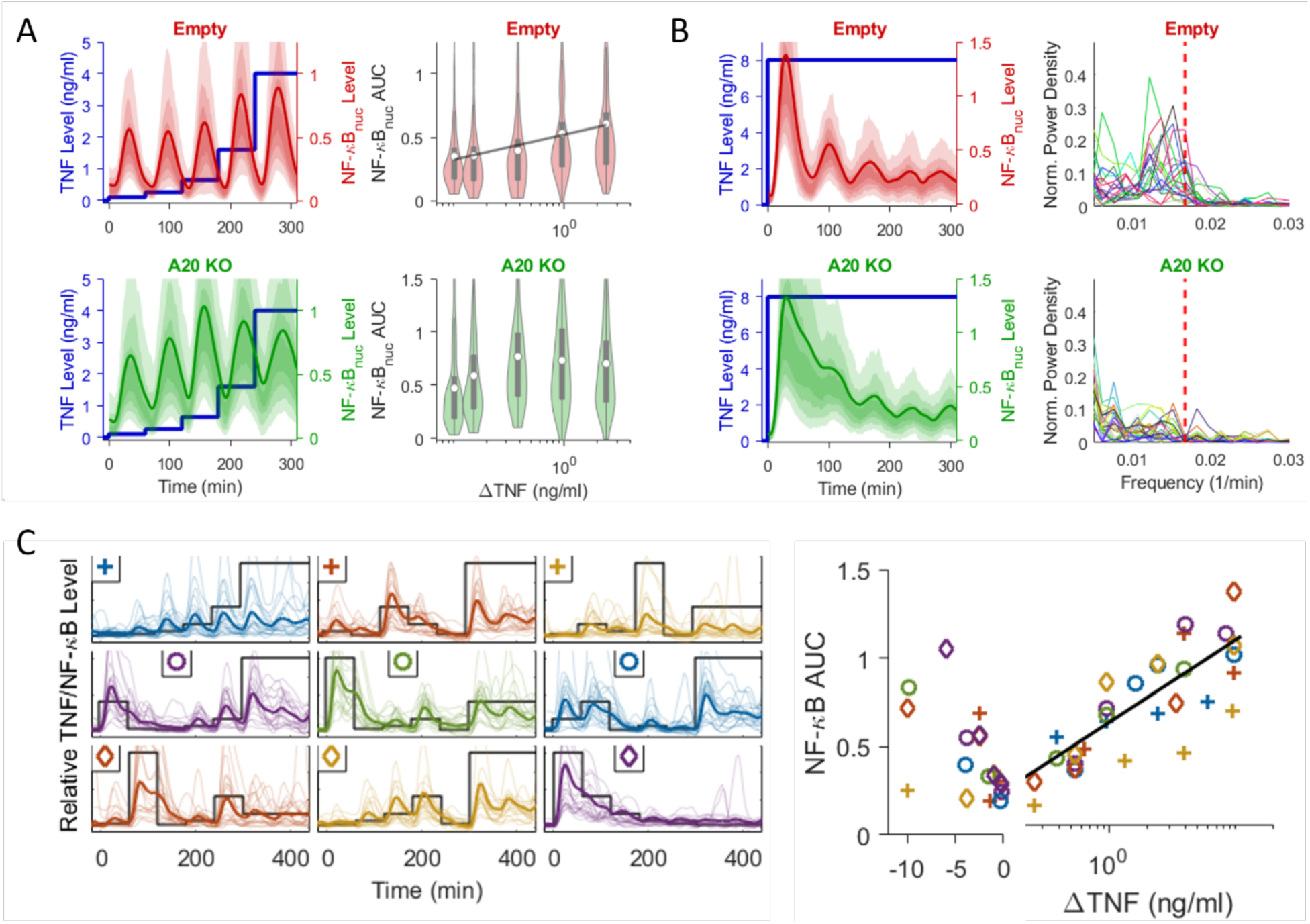
Measurements on A20 knock-out cells reveals the importance of A20 in sustained input differentiation. (A, B) To confirm the change in the NF-κB dynamics by the A20 deletion, A20 knock-out cells were generated (A20 KO) and NF-κB translocation measured under exponential TNF ramping (A) and instant increase (B). The results are compared to the wild-type background (Empty Vector). The right violin plots in (A) show the AUC at each time interval, where the white circles indicate the mean of population. For the instant increase in (B), a Fourier transform was applied to each cell trace to evaluate the power density spectrum. (C) Similar to wild-type in Fig. 2C, A20 knock-out cells were exposed to doses of TNF fluctuating from 0 to 10 ng/ml in random order (black line). The thick colored line shows the mean NF-κB trace, and the thin lines show 20 random single cell traces. On the right scatter plot, AUC after each dose increase/decrease was calculated then plotted in linear scale for negative ΔTNF (change in TNF concentration) and log scale for positive ΔTNF. The black line indicates the best fit to the log function.

### A20 controls IKK cycling that transmits dose change information to NF-κB

To examine the role of IKK cycling and A20 mediated negative feedback in dose differentiation more thoroughly, we gradually reduced the production level of A20 in our simulations and studied the change in NF-κB response under different TNF input dynamics (Fig. 5). The TNF level was increased in a step-wise fashion, and nuclear NF-κB and active IKK levels were evaluated. For unperturbed A20 translation, simulations successfully produced NF-κB amplitude that corresponded to the change in TNF concentration (Fig. 5A). However, when the translation rate of A20 is reduced to a quarter of its natural rate, we found that NF-κB localization could not track with changes in TNF input, disrupting the differentiator behavior. In this system, it takes longer for IKK_a_ level to return to baseline (Fig. 5B, blue curves), decreasing the level of IKK_n_ available to respond when the next stimulus comes. Consequently, the IKK_a_ production in the next input increase cannot fully reflect the increase in the TNF input dose, resulting in a damped response to input change. This effect becomes more severe if A20 is removed completely (Fig. 4C), and NF-κB cannot accurately reflect the increases in TNF dose.

Hence, the deletion of A20 results in two key outcomes: loss of feedback from nuclear NF-κB which transmits information about the previous cytokine dose, and longer adaptation time for IKK cycling to return to the ready state to reflect the change in the dose. The first would result in stronger NF-κB response at lower doses of TNF and earlier saturation of the NF-κB response during input ramping. The second would elongate the oscillation period for both IKK conformation and NF-κB translocation cycling. These anticipated outcomes of A20 deletion were observed in our experiments with the A20 knock-out (KO) cell line (Fig. 6A). To further confirm the longer relaxation time in the A20 mutant, we stimulated both wild-type background (designated as ‘empty vector’, EV) and A20 KO cells with a constant dose of TNF (8 ng/ml) continuously. While EV cells completed the initial cycle within 60 min of the stimulation, the nuclear NF-κB level of A20 KO cells was still decreasing even after 120 min (Fig. 6B). To assess how significantly A20 affects the oscillatory behavior at the single cell level, we applied Fourier transformation to each trace to evaluate power spectrum (Fig. 6B). This would reveal the most prominent oscillatory frequency in each cell. When A20 is knocked out, the mean frequency of the oscillation is decreased by more than half and more power is allocated on the zero frequency (the non-oscillatory response), demonstrating disruption of oscillatory NF-κB behavior.

Both exponential ramping and constant dose experiments indicate the critical role of A20 in generating prompt response to input change. To further confirm this, we delivered different doses of TNF in random order to A20 KO cells (Fig. 6C), similar to what we did with wild-type cells (Fig. 2C). Not only did the response to input change became noisier (correlation coefficient ∼ 0.76, compared to ∼ 0.91 for the wild type), but a significant level of NF-κB still remained in the nucleus even after the input level decreased, indicating the system’s reduced sensitivity to input dynamics.

Finally, to confirm that the behavior of our simulated A20 KO mutant was not biased by initial fitting with the full NF-κB circuit (Fig. 5A), we obtained the experimental best fit from the A20 knockout model directly (Supplementary Fig. 5A). Even after fitting the model without A20 directly, the resulting nuclear NF-κB traces cannot reproduce the traces observed from the different ramping experiments. Likewise, the best fit without IKK cycling also failed to reproduce the experimental results (Supplementary Fig. 5B). Hence, both our theoretical and empirical studies demonstrate the importance of A20 in resetting the IKK dynamics and receptor levels, and show that multipoint negative feedback enables accurate assessment of the input dynamics to produce appropriate NF-κB responses.

### Dynamic cytokine inputs create distinct NF-κB target gene expression profiles

NF-κB controls the expression of many target genes involved in immunity. To study the implications of NF-κB dose differentiation mechanism on target gene expression, we performed gene expression measurements on cells stimulated with different TNF dynamics. To determine whether NF-κB target gene expression dynamics differ between cytokine dosing patterns, we increased TNF level instantaneously and exponentially to 10 ng/ml in the microfluidic device and retrieved cells at different time points during and after the TNF ramping was finished. The mRNA levels of NF-κB target genes were subsequently measured through quantitative reverse transcription PCR (RT-qPCR) and compared with each other (Fig. 7).

**Figure 7.**
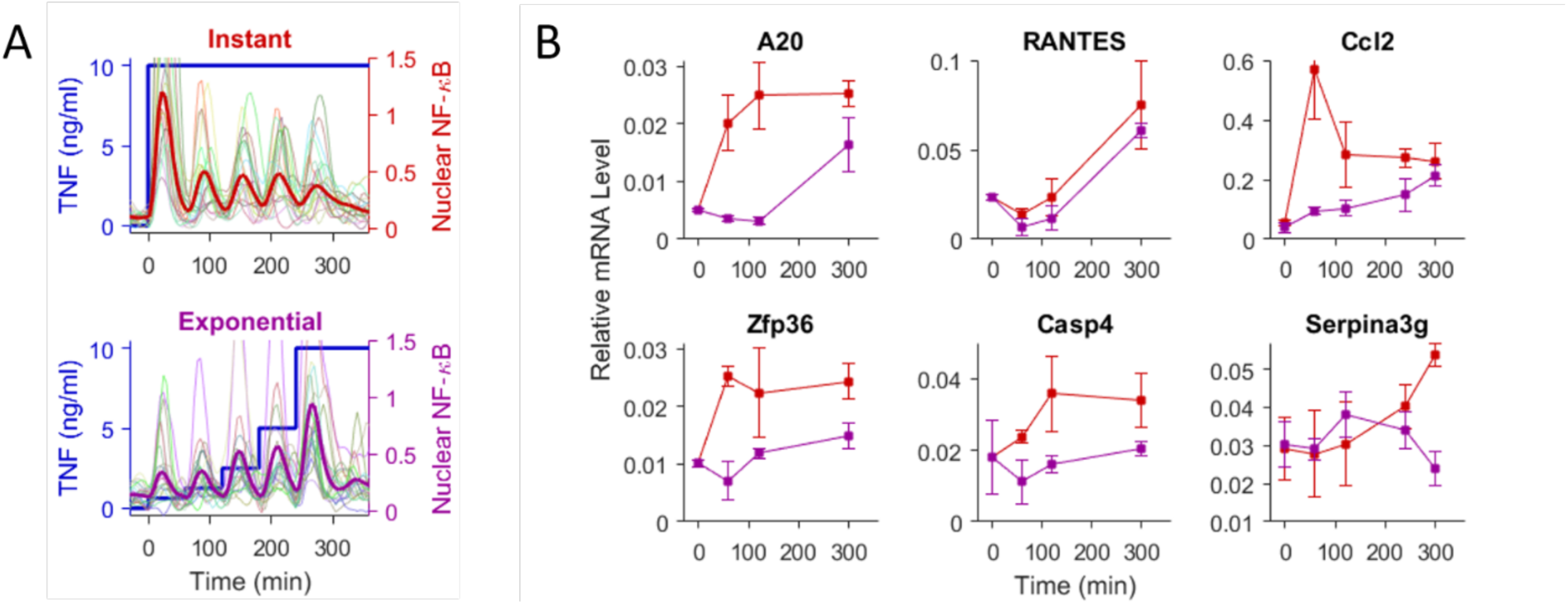
NF-κB target genes show distinct expression profiles when stimulated with different TNF dynamics. (A) Cells were stimulated with TNF in two different dynamics: instant increase to 10 ng/ml and exponential increase to 10 ng/ml (2-fold increase every hour for 5 h). To avoid any bias due to the TNF degradation, the chambers in the instant increase sample were also replenished with 10 ng/ml TNF medium every hour. (B) At multiple time points during the TNF ramping or constant feeding, cells were retrieved and immediately frozen for subsequent RT-qPCR measurement on NF-κB target genes. Red line shows the gene expression profile from instant increase, while the purple line shows the profile from the exponential ramping. The data points (squares) are the mean of two biological replicates, and error bars are the standard deviation from three technical replicates.

Surprisingly, we found that NF-κB target genes show remarkably different response patterns under dynamic cytokine input. Some genes closely followed the changes in cytokine concentration and NF-κB dynamic profiles, while others showed a preference to an instantaneous increase over a gradual increase in cytokine signal. For example, *A20* and *Ccl2* mRNA levels converged between the two dynamic patterns, with the instantaneous TNF increase resulting in a rapid increase in mRNA followed by maintenance of a high expression level, while mRNA levels under exponential TNF ramping gradually increased, following the NF-κB peak height. Notably, *Ccl2*, which recruits sentinel cells to the inflammation site^36^, also exhibited an initial sharp peak following instantaneous TNF increase; this pattern, which has been reported previously, may indicate complex temporal dynamics in downstream gene regulation^4^. In both cases, the mRNA levels in the exponential ramping caught up to those in the instant increase when the highest dose was reached. On the other hand, the expression of *Zfp36* and *Casp4*, which regulate inflammation and facilitate apoptosis^37–39^, exhibited minimal upregulation when TNF was increased exponentially, while an instantaneous increase caused a sharp increase in the expression level. These genes were largely unresponsive to a gradual increase in TNF concentration and required a large initial dose for significant activation.

*Serpina3g*, which prevents apoptosis^40^, exhibited gradual increase when the high TNF level persisted, but remained at the basal level when TNF concentration was increased exponentially. *RANTES*, another chemoattractant that recruits leukocytes, showed another distinct expression pattern. Regardless of instantaneous or exponential TNF increase, its expression level continuously increased, and the expression kinetics were almost indistinguishable between the input signal patterns.

## Discussion

We showed that during cytokine signaling, the NF-κB pathway accurately measures the changes in cytokine dose rather than the absolute dose itself, which is analogous to a differentiator circuit. Using microfluidic live-cell imaging experiments that enabled precise and continuous control of TNF and IL-1β concentrations in multitude of samples, we found that differentiator behavior is broadly applicable to various input dynamics, including instantaneous, linear and exponential increase, and random fluctuation. However, we found that NF-κB dose differentiation is not observed under bacterial PAM stimulation. This indicates that the regulation of NF-κB response significantly differs depending on the source of signaling molecules, and different response mechanisms are used for self-secreted cytokine signals and pathogenic signals such as TLR ligands.

We determined that multi-point negative feedbacks via A20 store short-term memory of the previous input level and reset the input transmission, consequently facilitating the differentiator behavior of NF-κB. Further, we demonstrated that the expression profile of NF-κB target genes changes significantly when different input dynamics are applied. Even though the final concentrations remained the same, some genes (*Zfp36, Casp4*, and *Serpina3g*) hardly showed any increase in their expression level during the increase in cytokine concentration (“input ramping”), while other genes’ expression (*A20* and *Ccl2*) showed close correspondence to the contemporary TNF dose during the ramping. These results suggest that cells actively tune their immune gene response according to the input dynamics.

NF-κB localization dynamics have been described in detail previously^18,41,42^. However, it is still unclear why signaling pathways employ these oscillatory responses to immune stimuli, which require complex circuits that are costly to the individual cell. Our study suggests that oscillatory response can transmit accurate information about the extracellular signaling input dynamics, i.e. how the concentration of the environmental signals change over time. An oscillating circuit involves time domain (frequency) and temporal change in the amplitude. Instead of relying on uncontrolled or natural degradation of signaling intermediaries, which can take from several hours or to days, the oscillating circuit can readily respond to the input change in the environment by actively degrading or resetting response regulators. Hence, oscillation itself could provide an optimized platform for measuring dynamic changes in the environmental inputs.

Previous *in vivo* and *in vitro* studies showed that during an inflammatory response TNF and IL-1β molecules are accumulated and degraded at different time scales, each exhibiting distinct kinetics^5,6,10^. How the NF-κB system would process the mixture of dynamic cytokines, is still unclear and can be complicated, especially when a distinct cytokine-specific feedback is involved in the signaling pathways^43^. Previous work has shown that A20 serves as the mediator for the signaling cross-talk between TNF and IL-1, as both stimuli strongly induced A20 expression and priming cells with IL-1 reduced their response to subsequent TNF^31^. This suggests that the differentiation behavior may not be restricted to the same stimulus (‘cis-differentiation’) but can carry across different stimuli (‘trans-differentiation’) with respect to the A20 controlled canonical pathway.

In conclusion, our work demonstrates the importance of cytokine dynamics for signal processing and downstream gene expression response in the NF-κB pathway. Cells employ multiple negative feedbacks and kinase recycling to repeatedly measure the dynamics of extracellular cytokine accurately. Depending on the dynamic pattern of cytokine increase (instant, linear, or exponential), cells regulate NF-κB target genes into different subsets, whose expression levels are increased or remained at the pre-stimulus level. Hence using any absolute cytokine level as the measure of the immune response or diagnosis of immune disease is not only flawed, but also may lead to ill-founded conclusion. Additionally, our single cell study revealed that the NF-κB response is strictly correlated to the rate of temporary input change. This is the first study that demonstrated a biological system can continuously and robustly differentiate the cytokine level in a consistent manner, regardless of widely varying dynamic signal profiles.

## Methods

### Cell culture

p65^-/-^ 3T3 mouse embryonic fibroblasts (MEFs) harboring p65-DsRed and H2B-GFP nucleus marker were cultured with DMEM (Gibco, 11965092) supplemented with 10% fetal bovine calf serum (HyClone™), 1% GlutaMax (Gibco), and 1× penicillin-streptomycin (Gibco) in regular flask. MEFs were maintained at 37°C and 5% CO_2_. Before reaching 100% confluency, cells were harvested using Trypsin-EDTA (Gibco) and diluted ∼1:10 in fresh medium for passaging every 2 or 3 days. A20 mutant cells (A20 KO) built based on this MEFs were also cultured and maintained in the same condition. For the experiment, FluoroBrite DMEM (Gibco, A1896702) with the same supplements was used to reduce the background fluorescence.

### Microfluidic device design

The previously designed and reported cell culture chip was modified for higher efficiency and success rate (Fig. 1C)^44^. The chamber size was increased to 3.5 x 0.8 mm to obtain more cells per chamber and images per sample, which improved image analysis and downstream gene expression data collection. The minimum distance between the features was adjusted to 150 μm for robust bonding and easier alignment. The input manifolds and flow pathways were revised for faster flushing between the sample feedings. Our new chip allowed more than 500 cells loaded per chamber and can run 64 independent samples (conditions) in parallel.

### Mold fabrication for microfluidic device

The master molds for the new design were fabricated by patterning photoresist deposited on silicon wafer through the multi-layer soft lithographic process^44,45^. Briefly, to fabricate the mold for the fluid layer, a positive photoresist (MicroChemicals, AZ40XT) was spin-coated on 4-inch wafer with spin rate 3400 RPM to get ∼ 25 μm feature height. After developing and hard-baking to get the rounded cross-sectional profile, the negative photoresist (MicroChemicals, SU-8 3025) was deposited on the top of the AZ layer by spin-coating at 3000 RPM, which produced ∼ 30 μm feature height. To fabricate the control layer, SU-8 3025 was spin-coated at 3500 RPM for ∼ 25 μm feature height. All exposure of photo-resist was done with maskless aligner (Heidelberg MLA150) using 375 nm laser and a dose of 280 mJ/cm^2^.

### PDMS microfluidic chip fabrication

The two-layer microfluidic chip was made by pouring and curing polydimethylsiloxane (PDMS) (Momentive, RTV-615) on these molds, and bonding control and flow layers. PDMS (66g, 10:1 monomer:catalyst) was poured onto the wafer patterned with flow layer design, air bubbles were removed under vacuum, and the PDMS was cured at 80 °C for > 5 h to make ∼ 2 cm thick PDMS slab with flow pattern grooved on the bottom. For the control layer, 10 g of PDMS was spun onto the wafer with control pattern at 2200 rpm to achieve ∼ 50 μm thickness, which is then cured at 80 °C for at least 1 h. After curing, the flow layer inputs were punched. Both PDMS layers were treated with oxygen plasma (Harrick, PDC-001), then aligned using a custom stereomicroscope. The aligned chip was cured at 80°C overnight to allow robust bonding between two layers. Next day, the holes to actuate the control layer were punched, then the chip was bonded to a glass slide (75 x 25 x 1 mm) through the same oxygen plasma treatment and baking procedure. Further details of the fabrication process can be found in our previous publication^46^.

### Microfluidic experiment setup

Before the experiment day, the valves in the control-layer of the microfluidic chip were connected to the pneumatic solenoid valves and electronic controller boxes. By actuating different sets of valves, the flow pathway in the microfluidic device could be manipulated either manually or automatically using custom-developed graphic-user-interface (GUI)^46^. Cell chambers in the chip were filled with a 1:3 fibronectin (1 mg/mL) in sterile PBS solution (Sigma Aldrich, FC010), and incubated overnight at room temperature for thorough coating. On the experiment day, chambers and channels were flushed with fresh medium, then the microscope box and brick (life imaging services GmbH) that enclose the microfluidic device were set at optimal environment for cell experiment (37°C, 5% CO_2_ level, and close to 100% humidity). Cells were concentrated to 5 × 10^6^ cells per ml to aim roughly 50% confluency when loaded into the microfluidic chambers., After loading cells, the chambers were closed for at least 1 h to allow cells to settle firmly before feeding any medium.

### High-throughput automated experiment procedure

The control layer of the microfluidic chip was actuated at 25 – 30 psi for thorough closure of the valves. The opening and closing of the valves were controlled via a custom graphic user interface (GUI) built with Matlab^46^. The GUI can automate the experimental procedure by running custom script, which greatly improved the accuracy, throughput, and reproducibility of the experiment. For each experiment, a new experiment code was written to direct the feeding timing and type of medium fed for each chamber. The media containing different concentrations of TNF-α (Life Technologies, PMC3014), IL-1β (R&D Systems™, 401ML010CF), and Pam2CSK4 (InvivoGen, tlrl-pm2s), were prepared right before the experiment and kept on ice during the experiment. All input media were delivered into the chip via polyetheretherketone (PEEK) tubing (VICI^®^, TPK.505) by pressurizing the vials containing the media. Input pressure was minimized (3 – 4 psi) to prevent shear stress on cells during feeding.

### Live-cell image acquisition

Phase contrast and epifluorescence images were acquired by Nikon Ti2 microscope enclosed within an incubator system (life imaging services GmbH). Images were captured with 20X magnification through CMOS camera (Hamamatsu, ORCA-Flash4.0 V2), and were collected every 5 min. For the p65-DsRed imaging, positions were imaged with 555 nm excitation with 0.5 – 1 sec exposure time and 75 – 100% LED intensity (Lumencor, Spectra X), while H2B-GFP was imaged under 485 nm excitation light with 50 – 100 msec exposure time and 25 – 50% LED intensity were used for nucleus marker (H2B-GFP). We did not notice any sign of photo bleaching even after collecting 150 sets of fluorescence images (equivalent to 12 h duration experiment). After each experiment, background fluorescence and dark frame images were taken for the flat field correction.

### Image analysis

NF-κB response or p65 translocation was assessed by analyzing fluorescence images with custom developed software (Matlab). The code first defines the position and boundary of each nucleus using H2B-GFP images. Combining this information across a sequence of images, the trajectory of each nucleus was evaluated^47^, which was then combined with p65-DsRed images to get single cell NF-κB traces. The median nuclear fluorescence was calculated to assess the NF-κB response in each cell. To quantify the background fluorescence for each cell *in situ*, the 20^th^ percentile of pixel brightness in a small sub-image encompassing each cell was evaluated, which was then subtracted from the corresponding p65 measurement. The resulting trace was smoothed using “lowess” method with span size of 5, in order to reduce the noise from cell movement, collisions between cells, and slight changes in the imaging focus. Using custom-developed analysis software (Matlab), each cell went through manual inspection to sort out cell death, cell division, or other defects that might affect the data quality. In this study, we inspected ∼ 100,000 cells manually, of which ∼ 30,000 were selected for subsequent data analysis.**Mathematical Modeling of NF-κB**

To capture the essential mechanism for the differentiation behavior in NF-κB pathway, a few approximations were applied to the mathematical analysis from our previous work^4^. The details of the mathematical modeling and fitting can be found in the Mathematical Method in Supplementary Information.

### CRISPR/Cas9-Gene editing of 3T3 cells

A20 gene in p65^-/-^ MEFs harboring p65-DsRed and H2B-GFP was edited through CRISPR/Cas9. A20-targeting guide RNAs (5’ – CACCGTTTGCTACGACACTCGGAAC – 3’, and 5’ – CACCGCTCGGAACTTTAAATTCCGC – 3’) were cloned into lentiCRISPR v2 (Addgene Plasmid #52961)^48^ and used for lentivirus production in HEK293T cells. After two days viral supernatants were collected and concentrated. Infected 3T3 cells were selected with Puromycin (2 µg/ml) for 3-5 days until cell death subsided. Knockout efficiency was assessed via Western blot (anti-A20: sc-166692, Santa Cruz) (Supplementary Fig. 6).

### Cell Retrieval from Microfluidic Device for RT-qPCR analysis

Before the automated experiment was finished, the corner of the microfluidic chip was cut open so that the outlet channel is exposed to air. When finished, cells in the chamber were trypsinized for ∼ 1 min to detach them from surface, then were directed to the open outlet channel by washing the chamber with phosphate buffered saline (PBS). The cutting allowed cells to accumulate at the end of outlet channel as a tiny droplet (2 – 3 μl), which can be retrieved easily using a small pipette. The retrieved droplet containing cells were immediately injected into the qPCR tubes containing 5 μl of lysis buffer (provided in CellsDirect Kit), and frozen at -80°C for subsequent NF-κB downstream gene measurements.

### RT-qPCR measurement

Cell lysis, RT, and pre-amplification done using CellsDirect One-Step RT-qPCR kit (ThermoFisher) as previously reported^46^. RT-qPCR run with custom (Supplementary Table 3) and predesigned TaqMan probes (Cat. No. 4453320 for *Ccl2*, and Cat. No. 4448892 for *Serpina3g*) on a Biorad CFX384 machine. Cq values calculated using software defaults and normalized to GAPDH. Relative expression levels calculated as 2^(Cq_Target_ – Cq_GAPDH_).

## Acknowledgements

We are grateful to Navid Ghorashian, Luke Vistain, Sara Saheb Kashaf, and Roy Wollman for valuable discussions and suggestions. M.O.M. was supported by a fellowship from the German Research Foundation (D.F.G.). This work was supported by NIH National Institute of General Medical Sciences (NIGMS) (R01GM117134-01) to A.H. and S.T.

## Author contributions

M.S., A.W., and H.T. did microfluidic experiments with help from P.P. and J.L. M.S and H.T. analyzed the experimental data with help from A.W. M.S. performed simulations with help from K.H. and A.M. M.O.M and A.W. constructed the A20 knockout cell line. A.W. and M.S. performed the RT-qPCR for downstream genes. S.T. supervised the work. M.S. wrote the initial draft. All authors contributed to the final version of the manuscript.

