## supplementary information for "Input dose differentiation by NF-κB"

for

**Son et al.**

**Table of Contents**

- **Supplementary Figure 1. TNF linear increase with various ramping rates shows robust TNF differentiation in the NF-κB system.**
- **Supplementary Figure 2. TNF exponential increase with various growth rates shows robust TNF differentiation in the NF-κB system.**
- **Supplementary Figure 3. Additional IL-1β ramping and random fluctuation experiments show IL-1β differentiation by the NF-κB system.**
- **Supplementary Figure 4. The differentiator behavior of the NF-κB system is sensitive to negative feedback loop parameters.**
- **Supplementary Figure 5. Fitting without A20 or the IKK module shows both are required for sustained differentiation.**
- **Supplementary Figure 6. Production of an A20 KO cell line.**
- **Mathematical Modeling of NF-κB control**
- **Supplementary Table 1. TNF dose and critical points, used for fitting.**
- **Supplementary Table 2. Parameters used for the simulation.**
- **Supplementary Table 3. List of all custom designed primers and probes used for qPCR**

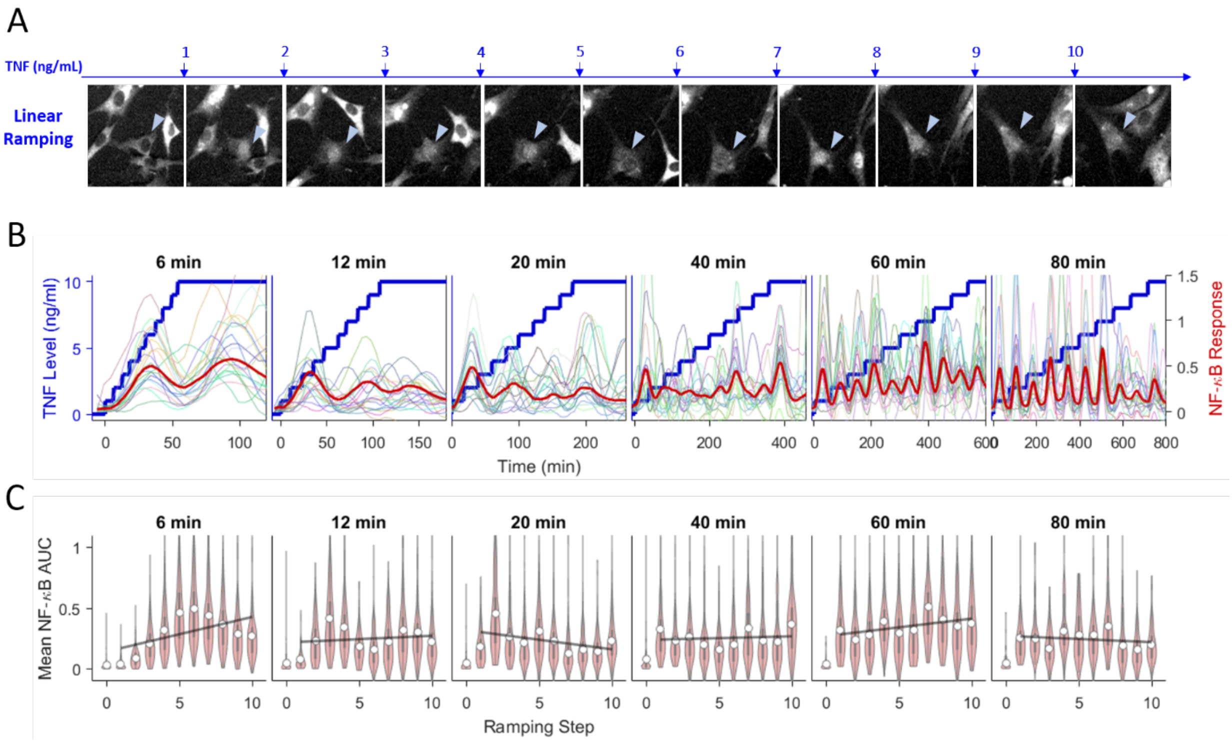


**Supplementary Figure 1. TNF linear increase with various ramping rates shows robust TNF differentiation in the NF-κB system.** Using microfluidics, the dose of TNF was increased by 1 ng/ml at different time interval until it reaches 10 ng/ml. Our results suggest that, except for the very fast ramping (time interval less than 12 min), the peak height or AUC for each TNF dose is maintained a roughly constant level during the various ramping rates. (A) NF-κB fluorescence images of a single cell in the 60 min interval ramping, showing an example of the constant activation. (B, C) Various ramping rates ranging from 0.75 ng/ml/h (80 min interval) to 10 ng/ml/h (6 min interval), were tested and plotted. In (B), thick blue line indicates the TNF dose change, thin colored lines represent 20 examples of the single cell traces, and thick red line shows the mean trace in the population (n > 200). For each ramping experiment, the area-under-curve (AUC) for each time interval was calculated and plotted in (C). Here, white circles show the mean of population, and black thick lines show the best fit = line.

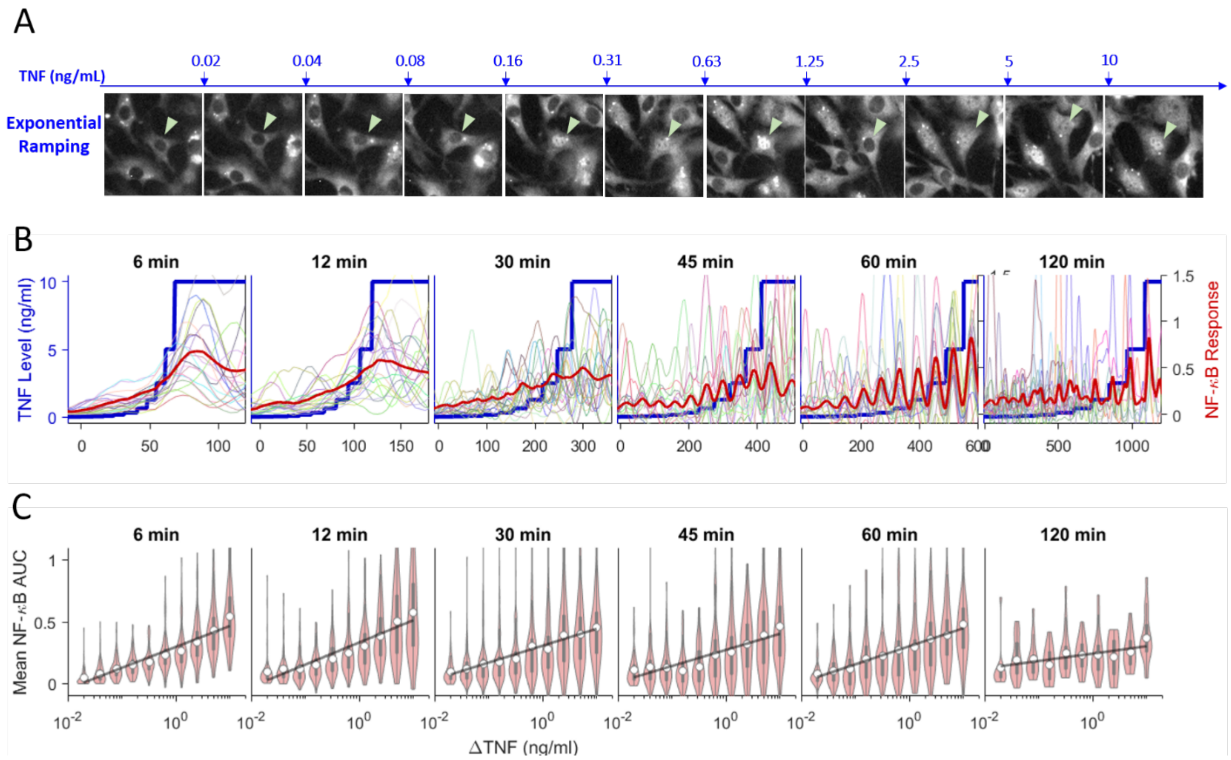


**Supplementary Figure 2. TNF exponential increase with various growth rates shows robust TNF differentiation in the NF-κB system.** The dose of TNF was doubled at different time interval until it reaches 10 ng/ml. For all doubling times tested, the peak height and AUC increased linearly during exponential ramping. (A) NF-κB fluorescence images of a single cell in the 60 min experiment, showing an example of the increasing NF-κB response during exponential ramping. (B, C) Various exponential growth constants ranging from 0.35/h (120 min interval) to 6.93/h (6 min interval), were tested and plotted. In (B), thick blue line indicates the TNF dose change, thin colored lines represent 20 examples of the single cell traces, and thick red line shows the mean trace in the population (n > 200). For each ramping experiment, the area-under-curve (AUC) for each time interval was calculated and plotted in (C). White circles show the mean step AUC of each population, and thick black lines show the best fit line to the log-differentiator behavior.

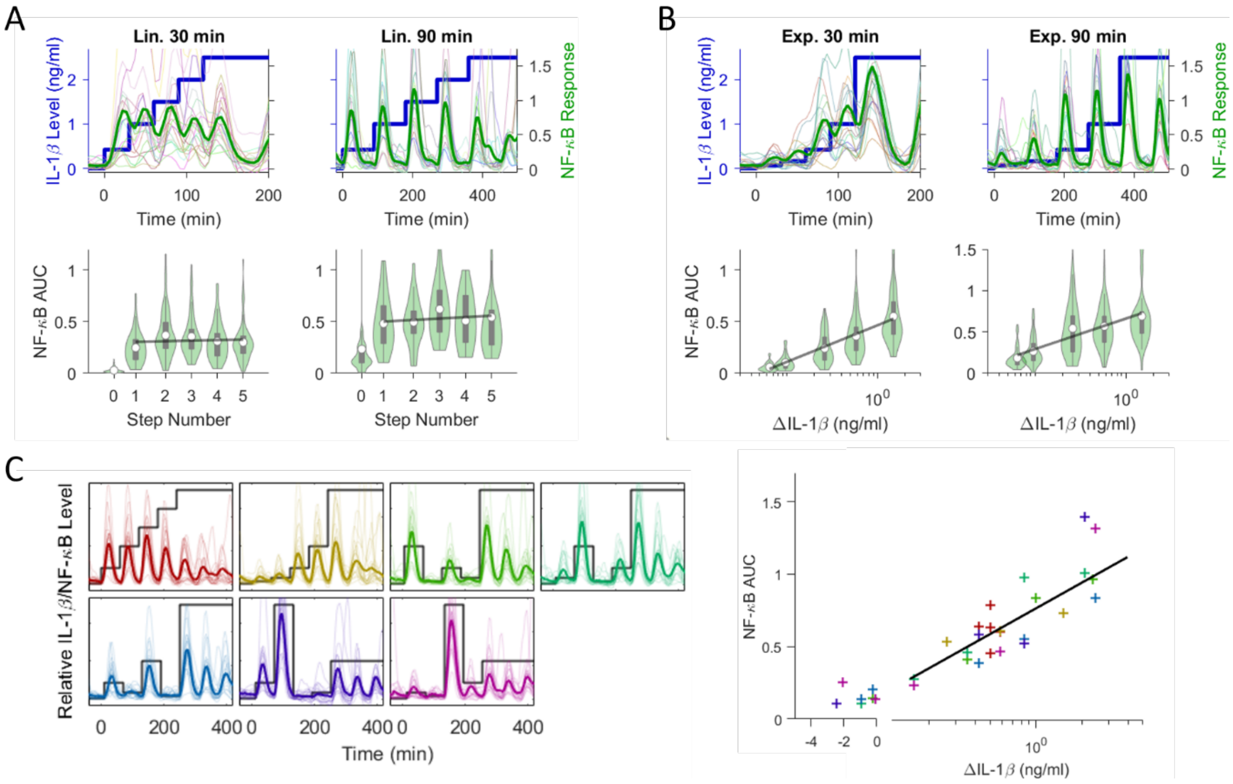


**Supplementary Figure 3. Additional IL-1β ramping and random fluctuation experiments show IL-1β differentiation by the NF-κB system.** (A, B) In addition to 60 min ramping (Fig. 2), 30 min and 90 min time intervals were also tested for IL-1β. For both linear and exponential ramping, the results are analogous to the differentiator behavior observed with TNF ramping. The peak height or AUC was maintained in linear ramping, while it increased linearly during exponential ramping. (C) Similar to the random TNF fluctuation experiment (Fig. 2C), cells were exposed to different doses of IL-1β in random order. The black line shows the IL-1β fluctuation ranging from 0 to 2.5 ng/ml, the thick colored line shows the mean NF-κB trace, and the thin lines show 20 random single cell traces for each sample. On the right scatter plot, the AUC after each dose increase/decrease was calculated then plotted in linear scale for negative IL-1β change and log scale for the positive IL-1β change. The black line indicates the best fit to the log function (correlation coefficient ~ 0.86).

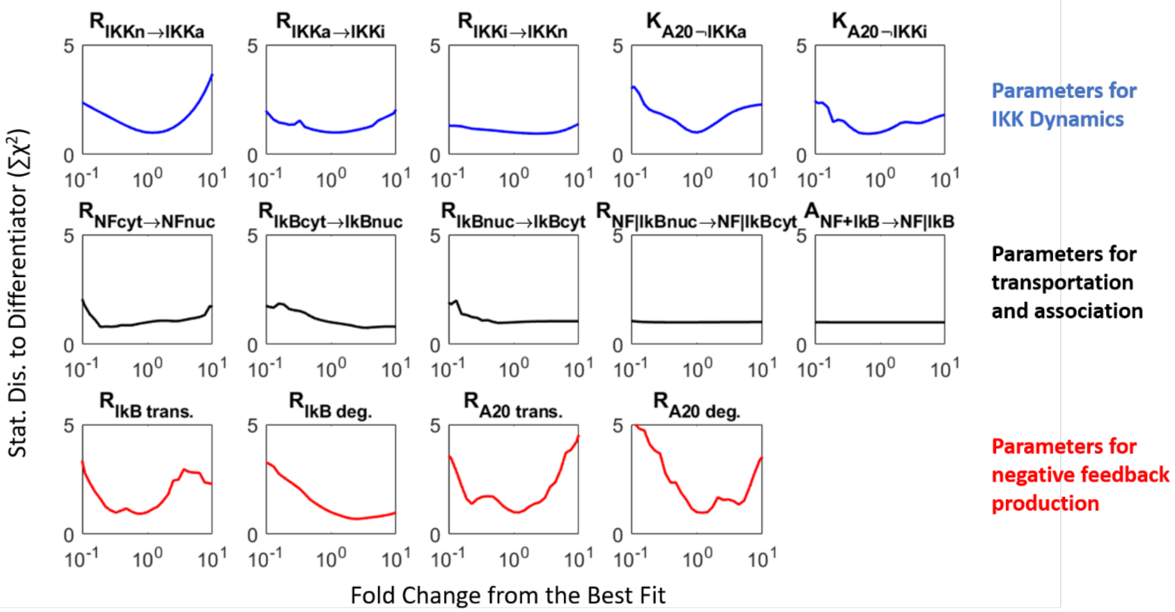


**Supplementary Figure 4. The differentiator behavior of the NF-κB system is sensitive to negative feedback loop parameters.** Parameter values involved in NF-κB regulation were varied 10-fold. The importance of each parameter for reproducing differentiation behavior in instant, linear, and exponential ligand increase was determined by simulating the NF-κB response under parameter variation and calculating the statistical distance ($\chi^2$) to the experimental mean response (Fig. 1 and 2). Here, the blue lines indicate the parameters involved in the regulation of the IKK cycling. From left to right, the parameters represent IKK activation rate by active TNFR, IKK inactivation rate, IKK neutralization rate, the dissociation constant for A20 inhibiting IKK activation, and the dissociation constant for A20 facilitating IKK inactivation. The red lines indicate the parameters directly related to the negative feedback from nuclear NF-κB (IκBα translation rate, IκBα degradation rate (including ones bound to NF-κB), A20 translation rate, and A20 degradation rate). The black lines represent other parameters associated with the transport of NF-κB and IκBα through the nuclear membrane, except the last one which indicates the association rate between NF-κB and IκBα. The full details of the modeling and simulation can be found in the Supplemental Information for the Mathematical Analysis.

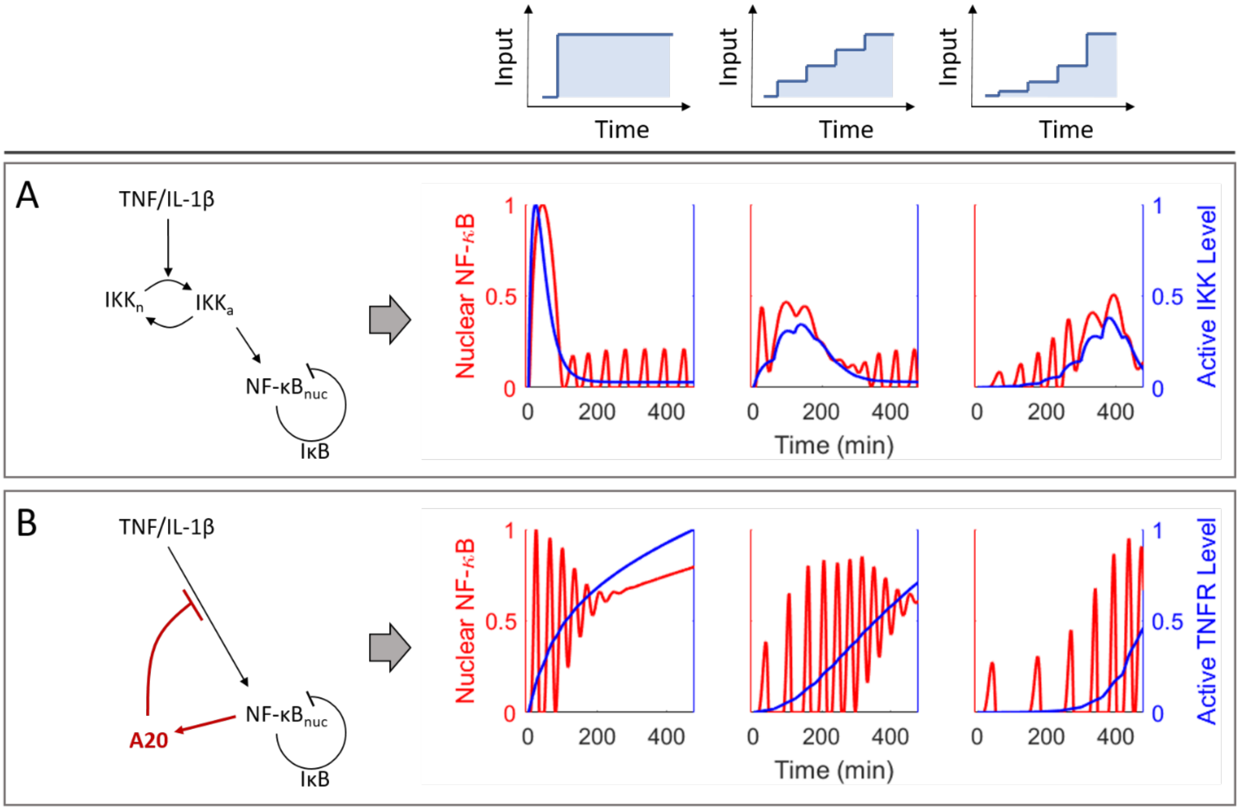


**Supplementary Figure 5. Fitting without A20 or the IKK module shows both are required for sustained differentiation.** (**A**) Using the same fitting method (Mathematical Method in Appendix), the NF-κB circuit without A20 was fit to the observed mean trace for each ramping pattern. Even after optimization, the NF-κB circuit lacking A20 failed to reproduce the differentiation behavior. The level of active IKK remained high during most part of the ramping resulting in the high trough level in the NF-κB oscillation. (**B**) Similarly, fitting was performed without IKK cycling to evaluate its importance. Here, we assumed the active cytokine receptors can directly degrade IκBα, and A20 can inhibit this process. Again, the best fit without IKK module failed to capture the differentiation behavior in the NF-κB oscillation.

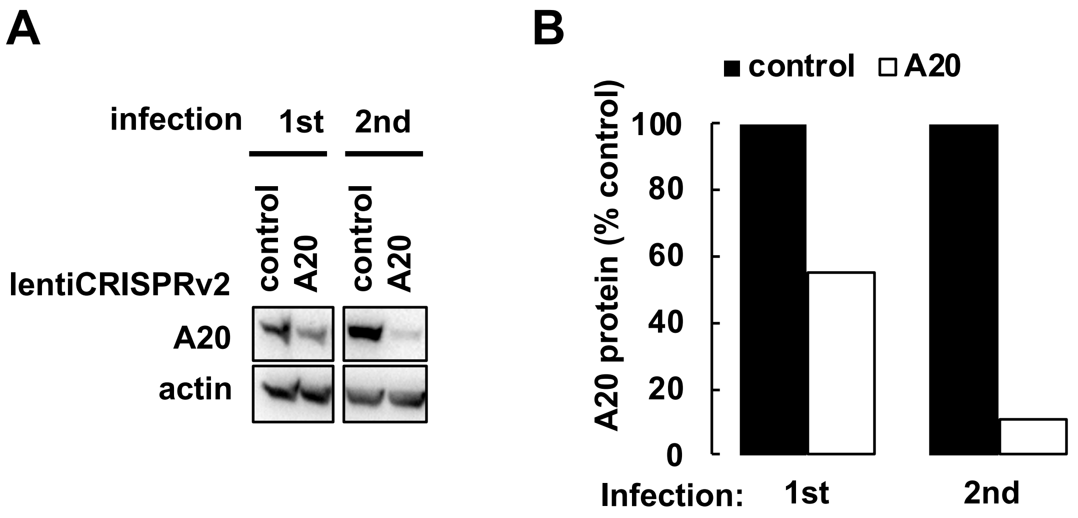


**Supplementary Figure 6. Production of an A20 KO cell line.** (A) Iterative rounds of CRISPR decreased levels of A20 protein in O1 3T3 cells by WB. (B) Quantification of relative intensity of A20 band between control and A20-targeted cells after each round of CRISPR.

**Mathematical Modeling of NF-κB control**

To capture the essential mechanism for the differentiation behavior in NF-κB pathway, a few approximations were applied to the mathematical analysis from our previous work [4]. The following three describe the changes.

1) The interactions between TNF, TNF receptor, and other intermediate IKK kinases (RIP1 and TRAFs) are summarized into one Hill function, whose dissociation constant and cooperativity were estimated based on the previous result [4].
2) Intermediate states of proteins that show only spontaneous kinetics once produced, are omitted. For example, the additional inactive state in IKK cycling after the initial inactivation (IKKi → IKKii → IKKn) is omitted (IKKi → IKKn). The phosphorylation of IκBα and its subsequent degradation (IκBα → IκBα-phosphorylated by IKKa → degraded), are simply expressed as IκBα being degraded by IKKa (IκBα → degradation by IKKa).
3) The active form of A20 is assigned to the dimer. A previous study demonstrated that A20 has multiple zinc finger binding motifs and its ubiquitin-binding functions occur in higher-order complexes [49]. Other works demonstrated that one of the motif (Zinc Finger 7) binds to the linear ubiquitin chain linked to active IKKγ (designated as NEMO in some studies) [50–53].

With these and a few minor simplifications, the number of system equations to build the model was reduced to ten compared to twenty in the previous model, and the number of parameters were reduced to twenty-five instead of forty-four. Since we only compared the simulated result to the mean of population response, we only considered deterministic reaction rates and built the ordinary differential equations (ODEs) to describe each species. The following describes the kinetics of each species in our simplified model. The full description of the parameters and their values are listed in Supplementary Table 2 at the end of this section.

**IKK in active state (*IKK$_a$*):** The first term describes the activation or transformation rate of the neutral IKK into active IKK through the phosphorylation by TNF. The IKK in its neutral state is expressed in terms of total IKK, amount of active IKK and inactivate IKK ($IKK_n = IKK_{Total} - IKK_a - IKK_i$). The intermediate process including TNFR activation and IKK kinase ubiquitination is summarized into a Hill function with only TNF. This activation is also attenuated by A20 dimers [29,30,49]. The second term describes inactivation rate (IKKa → IKKi), which is also mediated by A20 dimers [50–53]. Without A20, the conversion from $IKK_a$ to $IKK_i$ takes place at a slower rate [4].

(1) $\frac{d}{dt}(IKK_a)(t) = k_{IKKa} \times \frac{TNF(t)^{n_2}}{TNF(t)^{n_2}+(K_{TNF})^{n_2}} \times \left(IKK_{Total} - IKK_a(t) - IKK_i(t)\right) \times \frac{K1_{A20}}{A20_{dim}+K1_{A20}} - k_{IKKi} \times IKK_a(t) \times \frac{A20_{dim}+K2_{A20}}{K2_{A20}}$

**IKK in inactive state (*$IKK_i$*):** The first term describes the inactivation rate of $IKK_a$, which can be facilitated by $A20_{dim}$. The second term represents the dephosphorylation of IKKi to become neutral again.

(2) $\frac{d}{dt}(IKK_i)(t) = k_{IKKi} \times IKK_a(t) \times \frac{A20_{dim}+K2_{A20}}{K2_{A20}} - k_{IKKn} \times IKK_i(t)$

**Free NF-κB in cytosol (*$NFkB_{cyt}$*):** The first term describes the importation of cytosolic NF-κB into the nucleus. The second term describes the NF-κB|IκBα complex formation in the cytosol. The last two terms describe the degradation of IκBα in the cytosolic NF-κB|IκBα complex due to the catalytic phosphorylation by $IKK_a$ and due to the constitutive degradation. For simplification, the intermediate state for phosphorylated IκBα before the degradation is omitted. Hence, the rate term ($k_{IkB}$) for $IKK_a$ controlled degradation combines the association rate and degradation rate.

(3) $\frac{d}{dt}\left(NFkB_{cyt}\right)(t) = -I_{NFkB} \times NFkB_{cyt}(t) - A \times NFkB_{cyt}(t) \times IkB_{cyt}(t) + k_{IkB} \times IKK_a(t) \times \left(NFkB\middle|IkB_{cyt}\right)(t) + D_{IkB} \times (NFkB|IkB_{cyt})(t)$

**Free NF-κB in nucleus (*$NFkB_{nuc}$*):** The first term describes the importation of cytosol NF-κB into the nucleus, while the second term describes the depletion of nuclear NF-κB due to its association with nuclear IκBα. The same association rate was applied for both nuclear and cytosolic NF-κB|IκBα complex formation. However, the association in the nucleus is enhanced due to the smaller volume of the nucleus. The importation of NF-κB or IκBα into nucleus increases the concentration of each by the volume ratio.

(4) $\frac{d}{dt}(NFkB_{nuc})(t) = I_{NFkB} \times NFkB_{cyt}(t) - A \times V_{ratio}^2 \times NFkB_{nuc}(t) \times IkB_{nuc}(t)$

**mRNA transcription and degradation:** The first term describes the constitutive transcription, while the second term describes transcription induced by nuclear NF-κB. The last term describes the degradation rate. For simplification, our model uses the same mRNA term for both A20 and IκBα production.

(5) $\frac{d}{dt}(mRNA)(t) = C_{mRNA} + R_{mRNA} \times \frac{(NFkB_{nuc}(t))^{n_1}}{(NFkB_{nuc}(t))^{n_1}+(K_{mRNA})^{n_1}} - D_{mRNA} \times mRNA(t)$

**Free nuclear IκBα (*$IkB_{nuc}$*):** The first term describes the NF-κB|IκBα complex formation in nucleus. The second and third term describe the importation and exportation of free IκBα into and out of nucleus.

(6) $\frac{d}{dt}(IkB_{nuc})(t) = -A \times V_{ratio}^2 \times NFkB_{nuc}(t) \times IkB_{nuc}(t) + I_{IkB} \times IkB_{cyt}(t) - E_{IkB} \times IkB_{nuc}(t)$

**Free cytosolic IκBα (*$IkB_{cyt}$*):** The first term describes the NF-κB|IκBα complex formation in the cytosol. The second term accounts for the degradation of IκBα due to phosphorylation by $IKK_a$. The third term describes the translation IκBα protein from mRNA. The fourth and fifth term describe the importation and exportation of free IκBα into and out of the nucleus, while the last term describes the spontaneous degradation.

(7) $\frac{d}{dt}\left(IkB_{cyt}\right)(t) = -A \times NFkB_{cyt}(t) \times IkB_{cyt}(t) - k_{IkB} \times IKK_a(t) \times IkB_{cyt}(t) + R_{IkB} \times mRNA(t) - I_{IkB} \times IkB_{cyt}(t) + E_{IkB} \times IkB_{nuc}(t) - D_{IkB} \times IkB_{cyt}(t)$

**NF-κB|IκBα complex in cytosol (*NFkB|IkB$_{cyt}$*):** The first term describes the NF-κB|IκBα complex formation in the cytosol. The second and third term represent the release of NF-κB by the degradation of IκBα in the complex, one due to phosphorylation by $IKK_a$ and the other due to spontaneous degradation. The last term represents exportation of the complex from the nucleus.

(8) $\frac{d}{dt}\left(NFkB\middle|IkB_{cyt}\right)(t) = A \times NFkB_{cyt}(t) \times IkB_{cyt}(t) - k_{IkB} \times IKK_a(t) \times \left(NFkB\middle|IkB_{cyt}\right)(t) - D_{IkB} \times (NFkB|IkB_{cyt})(t) + E_{NFkB|IkB} \times (NFkB|IkB_{nuc})(t)$

**NF-κB|IκBα complex in nucleus (*NFkB|IkB$_{nuc}$*):** The first term describes the association of the NF-κB|IκBα complex in nucleus. The second term represents the exportation of the complex from the nucleus.

(9) $\frac{d}{dt}(NFkB|IkB_{nuc})(t) = A \times V_{ratio}^2 \times NFkB_{nuc}(t) \times IkB_{nuc}(t) - E_{NFkB|IkB} \times (NFkB|IkB_{nuc})(t)$

**A20 protein and dimerization (*A20* and *A20$_{dim}$*):** The first term represents the translation of the A20 protein, while the second term describes the constitutive degradation. For the A20 dimer, steady-state approximation was applied for simplification. Note that eq. (11) can be integrated into eq. (1) and (2) by substitution, hence leaving total of ten ODEs.

(10) $\frac{d}{dt}(A20)(t) = R_{A20} \times mRNA(t) - D_{A20} \times A20(t)$

(11) $A20_{dim}(t) = K_{dim} \times A20(t)^2$

**Model Fitting and Parameter Evaluation**

The above ODEs include twenty-five parameters, of which ten were found or estimated from this and previous studies. The rest of parameters were fitted to our experimental results with various TNF ramping (Fig. 1 and 2). We highlight that the TNF degradation was not considered in order to show the robust differentiation by NF-κB system. Due to many approximations employed in the model, it is not feasible to reproduce the detailed kinetics of NF-κB response. This simulation aims to reproduce only the generic behavior of NF-κB response and the log-differentiation behavior of the system mediated by A20, IκBα, and IKK cycling. For this, we selected following pivotal points in the NF-κB oscillation: peak heights, peak times, and trough heights (Supplementary Table 1).

| Ramping Pattern | | 1st feeding | 2nd | 3rd | 4th | 5th | 6th | 7th | 8th |
|---|---|---|---|---|---|---|---|---|---|
| Instant | TNF (ng/ml) | 10 | 10 | 10 | 10 | 10 | 10 | 10 | 10 |
| | Rel. peak height | 1 | 0.4 | 0.3 | 0.2 | 0.2 | 0.2 | 0.2 | 0.2 |
| | Peak time (min) | 20 | 82 | 144 | 206 | 268 | 330 | 390 | 450 |
| | Rel. trough height | 0.05 | 0.05 | 0.05 | 0.05 | 0.05 | 0.05 | 0.05 | 0.05 |
| Linear | TNF (ng/ml) | 1 | 2 | 3 | 4 | 5 | 6 | 7 | 8 |
| | Rel. peak height | 0.45 | 0.45 | 0.45 | 0.45 | 0.45 | 0.45 | 0.45 | 0.45 |
| | Peak time (min) | 25 | 86 | 147 | 208 | 269 | 330 | 390 | 450 |
| | Rel. trough height | 0.05 | 0.05 | 0.05 | 0.05 | 0.05 | 0.05 | 0.05 | 0.05 |
| Exponential | TNF (ng/ml) | 0.08 | 0.16 | 0.32 | 0.64 | 1.25 | 2.5 | 5 | 10 |
| | Rel. peak height | 0.05 | 0.15 | 0.25 | 0.35 | 0.45 | 0.55 | 0.65 | 0.75 |
| | Peak time (min) | 30 | 90 | 150 | 210 | 270 | 330 | 390 | 450 |
| | Rel. trough height | 0.01 | 0.02 | 0.03 | 0.04 | 0.05 | 0.05 | 0.05 | 0.05 |

**Supplementary Table 1** TNF dose and critical points, used for fitting. The peak and trough heights were normalized to the maximum height, which happens at first peak for 10 ng/ml TNF instant ramping.

For the fitting, the statistical distance ($\chi^2$) to each of values listed in Supplementary Table 1 was calculated. The sum of all statistical distances (there are total of 48 values to fit) was used as metric to evaluate the goodness of fit. Each ramping pattern and each point were equally weighted, and the sum of statistical distances was minimized using "fminsearch" function in Matlab with practical boundary conditions (range) for the parameters. To ensure the best fit, we also used the globalsearch function in Matlab, which randomly selects multiple start parameters within the range and runs a minimization search. Supplementary Table 2 shows the parameter values found from fitting or evaluated based on previous studies.

| Parameter | Value | Description (reference) |
|---|---|---|
| $V_{ratio}$ | 4 | Ratio of cytoplasmic to nuclear volume. [54] |

| | | |
|---|---|---|
| $n_1$ | 3 | Cooperativity for the mRNA transcription by nuclear NF-κB. [32,55] |
| $NFkB_{total}$ | $3.5 \times 10^4$ | Total amount of NF-κB. [56] |
| $IKK_{total}$ | $6 \times 10^3$ | Total amount of IκB kinase. [56] |
| $K_{TNF}$ | 35 ng/ml | Dissociation constant for TNF activating TNF receptor and $IKK_n$ (estimated from [4]) |
| $n_2$ | 1.5 | Hill coefficient for TNF activating TNF receptor and $IKK_n$ (estimated from [4]) |
| $k_{IKKa}$ | 0.55 $min^{-1}$ | Maximum transformation rate from IKKn to IKKa (fitted) |
| $k_{IKKi}$ | 0.045 $min^{-1}$ | Transformation rate from IKKa to IKKi (fitted) |
| $k_{IKKn}$ | $4.5 \times 10^{-4}$ $min^{-1}$ | Transformation rate from IKKi to IKKn (fitted) |
| $I_{NFkB}$ | 0.057 $min^{-1}$ | Importation rate of cytoplasmic NF-κB into nucleus (fitted) |
| $I_{IkB}$ | 0.082 $min^{-1}$ | Importation rate of cytoplasmic IκBα into nucleus (fitted) |
| $E_{IkB}$ | 0.471 $min^{-1}$ | Exportation rate of nuclear IκBα into cytoplasm (fitted) |
| $E_{NFkB\|IkB}$ | 0.728 $min^{-1}$ | Exportation rate of nuclear NF-κB\|IκBα into cytoplasm (fitted) |
| A | 0.268 $min^{-1}$ | Association rate between NF-κB and IκBα (fitted) |
| $K_{mRNA}$ | 3227 | Dissociation constant for nuclear NF-κB inducing mRNA. The value is roughly 10% of total NF-κB (fitted) |
| $C_{mRNA}$ | 0.01 $min^{-1}$ | Constitutive transcription rate for mRNA (estimated from [4,57]) |
| $R_{mRNA}$ | 20 $min^{-1}$ | Maximum transcription rate induced by nuclear NF-κB (estimated from [4]) |
| $D_{mRNA}$ | 0.035 $min^{-1}$ | Degradation rate for mRNA (measured in this study) |
| $R_{IkB}$ | 60.7 $min^{-1}$ | Translation rate for IκBα (fitted) |
| $k_{IkB}$ | 0.0051 $min^{-1}$ | Degradation rate of IκBα mediated by $IKK_a$ (fitted) |
| $D_{IkB}$ | 0.0056 $min^{-1}$ | Spontaneous degradation rate of IκBα (fitted) |
| $R_{A20}$ | 10 $min^{-1}$ | Translation rate for A20 (assumed) |
| $D_{A20}$ | 0.108 $min^{-1}$ | Spontaneous degradation rate of A20 monomer (fitted) |
| $K1_{A20}/K_{dim}$ | $1.37 \times 10^8$ | The ratio between Michaelis coefficient in TNFR activity attenuation and association constant for A20 dimerization (fitted) |
| $K2_{A20}/K_{dim}$ | $1.37 \times 10^8$ | The ratio between Michaelis coefficient in IKKa inactivation and association constant for A20 dimerization (fitted) |

**Supplementary Table 2** Parameters used for the simulation. All substrates except TNF cytokine are quantified in numbers of molecules, not in concentration unit.

We note that even with many approximations used in the model, most of fitted parameters are comparable to the parameters from the previous studies [1,4,32], exhibiting less than 5-fold difference. The parameters that showed significant difference are the rate terms in IKK cycling (activation, inactivation, and neutralization), and A20 degradation rate. The different rates in IKK cycling are due to the smaller number of IKK molecule used in our model, $6x10^3$ instead of $2x10^5$. The number of NEMO molecules, an important subunit in IKK complex, has been measured to be around $6x10^3$ [56]. Also we did not include another inactive state of IKK (designated as $IKK_{ii}$ in other studies) for the simplification purpose. Hence, our neutralization rate term combines another inactivation rate and actual neutralization rate in other models. Given these modifications in our model, our values are in a comparable range to the values from other studies. The high degradation rate of A20 is due to the dimerization mechanism newly employed in this study [49]. We used the steady state approximation for the A20 dimer formation, where the dimer needs to disintegrate before degradation. Hence, in our model most of A20 exists in dimeric form, whose amount is proportional to the squared amount of A20 monomer. Missing other key elements in A20 high-order complex formation can be also the cause of high degradation rate in our fitting. For example, a previous study demonstrated that the active form of A20 requires binding to adaptor molecules including TAX1BP1, Itch, and RNF11 [30].

**QUANTIFICATION AND STATISTICAL ANALYSIS**

The method for statistical test, fitting, and plot are described in the corresponding Figure legends. All statistical tests including R-squared, $\chi^2$, Pearson correlation coefficient evaluation and clustering (Fig 3) were calculated using Matlab 2017b and its Statistics toolbox. Details for the statistical test for parameter sensitivity (Fig. 4 and Supplementary Fig. 4), are described in the Model Fitting and Parameter Evaluation section in the Mathematical Method. For the violin plots, we adapted a custom-developed function obtained at https://github.com/bastibe/Violinplot-Matlab.

| **Primers & Probes** | **Sequence** |
|---|---|
| A20 Forward | GCA GCT GGA ATC TCT GAA ATC T |
| A20 Reverse | AGT TGT CCC ATT CGT CAT TCC |
| A20 Probe | /56-FAM/AA ACA GGA C/ZEN/T TTG CTA CGA CAC TCG G/3IABkFQ/ |
| ZFP36 Forward | CCA TCT ACG AGA GCC TCC A |
| ZFP36 Reverse | TTT ATG TTC CAA AGT CCT CCG A |
| ZFP36 Probe | /56-FAM/AT GAG CCA T/ZEN/G ACC TGT CAT CCG AC/3IABkFQ/ |
| Casp4 Forward | AGG TGA AAT GCT TCT CCA GAC |
| Casp4 Reverse | TTC TTC TGG TTC CTC CAT TTC C |
| Casp4 Probe | /5Cy5/AT TCT TCA G/TAO/T GTG GAC CCA GGC AG/3IAbRQSp/ |
| RANTES Forward | GCC CAC GTC AAG GAG TAT TT |
| RANTES Reverse | CTT GAA CCC ACT TCT TCT CTG G |
| RANTES Probe | /5Cy3/TT TGT CAC TCG AAG GAA CCG CCA A/3IAbRQSp/ |

**Supplementary Table 3:** List of all custom designed primers and probes used for qPCR (A20, ZFP36, Casp4, RANTES). Primers and probes for the other genes, Serpina3g and Ccl2, were commercially available.
